# Post-stroke rapamycin treatment improves post-recanalization cerebral blood flow and outcome in rats

**DOI:** 10.1101/2023.11.16.567392

**Authors:** Anna M Schneider, Yvonne Couch, James Larkin, Alastair M Buchan, Daniel J Beard

**Affiliations:** Acute Stroke Programme, Radcliffe Department of Medicine, University of Oxford, Oxford, UK; Department of Oncology, University of Oxford, Oxford, UK; School of Biomedical Sciences and Pharmacy, University of Newcastle, Australia

**Author notes:** Corresponding author: Professor Alastair M. Buchan, DSc, FMedSci Acute Stroke Programme, Room 7501, Level 7A/B, John Radcliffe Hospital, Oxford OX3 9DU, United Kingdom. These authors contributed equally to this work (Alastair M Buchan, Daniel J Beard).

**Keywords:** focal ischemia, mTOR, rapamycin, neuroprotection, MRI

## Abstract

Ischaemic stroke treatment is limited to recanalizing the occluded vessel, while there is no approved adjunctive cerebroprotective therapy to protect either the neurons and parenchyma or the neurovascular unit. Pharmacological inhibition of mammalian target of rapamycin-1 (mTORC1) with rapamycin has shown promise in reducing infarct volume and improving functional outcomes. However, previous studies that investigated the effects of rapamycin on the vasculature and cerebral blood flow (CBF), administered rapamycin prior to or during stroke induction, thus limiting the potential for clinical translation. Therefore we investigated whether rapamycin maintains its cerebrovascular protective effect when administered immediately after recanalization following 90 minutes stroke in Wistar rats. We show, that rapamycin significantly improved post-recanalization cerebral blood flow (CBF), suggesting a beneficial neurovascular effect of rapamycin. Rats treated with rapamycin had smaller infarct volumes and improved functional outcomes compared to the control animals at three days post-stroke. The mechanisms of the overall positive effects seen in this study are likely due to rapamycin’s hyperacute effects on the neurovasculature, as shown with increased CBF during this phase. This paper shows that rapamycin treatment is a promising adjunct cerebroprotective therapy option for ischemic stroke.

## Introduction

Stroke treatment has recently been revolutionized by recanalization therapy and better patient selection thanks to improved imaging modalities. Despite high recanalization rates (60-70%), half of patients are still left with poor functional outcomes, and only about 10% are symptom-free at three months (1). Therefore, an adjunct pharmacological treatment that has neuronal and vascular protective effects may help improve outcomes post-recanalization.

Pharmacological inhibition of mTORC1 with the FDA-approved anti-rejection medication rapamycin has been shown to protect brain tissue from dying in experimental models of stroke (2, 3). However, the mechanism by which rapamycin exerts its cytoprotective effects and which cerebral cells and anatomical structures it targets are still being explored. A large body of literature reports that rapamycin can reduce infarct volume and improve neurological function in stroke models (4–19). However, only a few papers have investigated the effects of rapamycin treatment on structural changes of the blood-brain barrier (BBB) and cerebral blood flow (CBF) after transient middle cerebral artery occlusion (tMCAo) (2, 8, 9, 13, 14, 18, 19), of which five reported decreased BBB breakdown due to rapamycin treatment (9, 13, 14, 18, 19). Our recent work has highlighted that rapamycin can also enhance collateral and post-recanalization CBF (Beard et al. 2020. However, many of these studies either treated animals before the onset of stroke or prior to recanalisation. Therefore it is not clear if rapamycin maintains its cerebrovascular protective effect if administration is delayed to immediately after recanalization, thus better matching the clinical scenario of administration of an adjunct cerebroprotectant with vessel recanalization.

There has been a concerted effort from the preclinical stroke field to improve the quality and thus reduce the bias in clinical studies (20). While our recent meta-analysis of rapamycin showed that no studies have completely adhered to these guidelines, we have designed the present study in such a way that it avoids these shortcomings (2). In addition to the clinically translatable study design of administering rapamycin in the post-reperfusion phase, the methods used to assess the drug’s effect have also been chosen to be comparable to clinical stroke practice.

MRI has proven to be an optimal tool for the longitudinal assessment of cerebral blood flow (CBF), lesion progression and for evaluating translational therapeutic studies (21, 22). Sequences used in MRI can detect different aspects of the ischemic lesion (23), and because brain damage following cerebral ischemia is a dynamic process, assessment with MRI requires varied imaging modalities to suit the particular pathophysiologic state of the lesion. For example, T2w-MRI provides anatomical information, whereas more dynamic sequences such as diffusion- and perfusion-weighted imaging (DWI and PWI, respectively) can detect tissue microstructure, metabolism, and hemodynamics, and in stroke, inform about potentially salvageable brain tissue using the DWI/PWI mismatch coefficient (23). In this study, we used different MRI sequences at the three days post-stroke onset to study rapamycin’s effect on perfusion and lesion volume, while functional testing assessed the impact on behavioral outcomes and its correlation to the MRI data. The aims of this study are threefold. First, to understand how mTORC1 inhibition with rapamycin affects immediate post-recanalisation CBF. Second, to examine if the effects of rapamycin on perfusion changes persist out to 3 days post-stroke, and to determine any effects on lesion volume and functional outcomes. Third, to study the effect of mTORC1 inhibition on the integrity of the BBB using contrast agent MRI.

## Materials and methods

### Ethics and animal care

All experimental procedures were approved by the UK Home Office (1986 Animal Act, Scientific Procedures), conducted in accordance with the Clinical Medicine Ethical Review Guidelines of the University of Oxford, and conformed to the ARRIVE and IMPROVE guidelines for animal and pre-clinical stroke work (24, 25). Male Wistar Han rats (250g - 320g, 8 – 11 weeks old, Envigo Research Model Services in Blackthorn, England) were housed in individually ventilated cages under a 12-hour light/12-hour dark cycle with *ad libitum* access to food (standard food pellets) and water. The principal investigator began daily animal handling, weighting, and training the rats for the adhesive removal test (see below) 3 days before the surgery. During the 3 days following surgery, the rats underwent daily welfare scoring.

### Study design

#### Controls and exclusion criteria

Appropriate control groups were included in all experiments. For the comparison with rapamycin, vehicle groups were used. Since naïve groups were included in similar animal studies undergoing the same intervention, and the investigators tried to minimize the number of animals used, no sham groups were included in this study (9). Where possible, within-subject controls were used, for example, when comparing the extent of stroke expansion on the ipsilateral versus contralateral hemisphere. Pre-defined exclusion criteria were a laser Doppler flow (LDF) signal decrease of less than 70% from baseline and a confirmed subarachnoid haemorrhage (SAH), a rare but known complication of the focal ischemic stroke model used.

#### Randomization and blinding

Animals were randomized with the sealed envelope method to receive either rapamycin (Sigma Aldrich, 250 μg/kg, n=9) or vehicle (<5% ethanol in saline, n=9), which was administered by a surgeon blind to treatment allocation (AMS). Blinding was continued throughout the whole experiment and until data acquisition was complete.

### Anesthesia and Monitoring

Rats were initially anaesthetized by inhaling 5% isoflurane in 70% N_2_O and 30% O_2_ and maintained at 1-2% isoflurane in 70% N_2_O and 30% O_2_. During the surgery, the core body temperature of all animals was maintained at 37.0 ± 0.5°C using a rectal thermometer connected to a feedback-controlled heating pad (Harvard Apparatus, Cambourne, UK). Physical parameters, including body temperature and respiratory rate, were checked and recorded every 15 minutes throughout the surgical intervention. Respiration was kept between 50 and 60 breaths per minute by adjusting the isoflurane concentration.

### Cerebral blood flow measurements

A fibreoptic probe was firmly positioned on the intact skull surface based on locations described previously (26). An LDF apparatus (Oxyflo 2000 Optronix, Oxford, UK) was used to continuously monitor cerebral perfusion of the lateral MCA territory, corresponding to the core territory after MCA occlusion. LDF traces were recorded onto a Windows XP workstation running with WinDaq Data Acquisition Software (Dataq Instruments, Akron, Ohio, USA). MCAo was confirmed by a >70% decrease in LDF signal from baseline. Selection of a 70% decrease as the threshold was chosen based on previous studies reporting an infarct threshold of 20-30% from baseline (27, 28).

### Focal cerebral ischemia

This study used 22 rats. All surgical procedures were conducted under sterile conditions. Focal brain ischemia was induced by transient occlusion of the MCA for 90 minutes, according to a previously described method (29–31). Reperfusion was confirmed by visual reperfusion of the proximal ICA and a sudden increase of the recorded cerebral blood flow in the MCA territory. Immediately after reperfusion, rapamycin or vehicle was administered through intravenous tail vein injection. The CBF was recorded for another 90 minutes from the time of reperfusion.

### Behavioral assessment

Neurological assessments to measure neurological impairments post-stroke were performed on all three days following surgery. Tests involved Bederson (32), Garcia (33), and the adhesive removal test (34). The Bederson scale studies post-stroke behavioral deficits by assessing forelimb flexion, resistance to lateral push, and circling behavior, and using a grading scale of 0 to 3, with 0 being not affected and 3 being severely affected (32). The Garcia testing battery consists of 6 tests to evaluate sensorimotor deficits, where performance is graded on a scale of 0-3, with 0 being not affected and 3 being severely affected by the stroke, representing a minimum score of 0 and a maximum score of 18 points (33). The adhesive removal test involves placing a plaster on the forepaw and measuring the “time-to-contact” (somatosensory deficit in the contralesional forepaw) and “time-to-remove” (contralesional forepaw dexterity). To decrease inter-individual differences within this test, training sessions from 3 days before the surgical procedure (2 trials per day) were performed, and the time used for noticing and removing the sticky tape on the contralateral forepaw was analyzed and compared among the rapamycin and vehicle groups.

### Magnetic resonance imaging

All MRI acquisitions were carried out on a 9.4T horizontal-bore scanner (Agilent Technologies Inc., Santa Clara, USA) with a 72 mm volume transmit coil and a 4-channel surface receive array (Rapid Biomedical, Rimpar, Germany). The animal’s anesthesia was induced with 5% isoflurane in 70% N_2_ and 30% O_2_ and maintained at 1-2% isoflurane in 70% N_2_ and 30% O_2_. The rats were placed in a cradle equipped with a stereotaxic holder, a rectal thermometer, and a pressure probe to monitor respiration. The rectal thermometer was connected to a feedback-controlled heating pad (Harvard Apparatus, Holliston, United States) to maintain core body temperature at 37.0 ± 0.5°C, and respiration was kept between 50 and 60 breaths per minute by adjusting isoflurane concentration. Physical parameters were recorded and checked regularly throughout the imaging session.

Acquisition of baseline maps for T1 and T2 relaxation times (seconds), arterial spin labeling (ASL) for measuring cerebral blood flow (CBF) (mL/100g/min), and apparent diffusion coefficient (ADC) for delineating diffusion (μm²/ms) were performed. T1 and T2 data were acquired using a spin-echo echo-planar imaging (EPI) readout with FOV = 32 x 32 mm², matrix = 256 x 256 mm², thickness = 1 mm, 10 slices. T1w images were taken with a scan repetition time (TR) and echo time (TE) of 500 ms and 20 ms, respectively. The parameters were identical for imaging acquisition, both before and after administration of 150 μL gadodiamide (Omniscan, Germany) via an indwelling tail vein cannula. T1-map scans (repetition time/echo time [TR/TE] = 1000 ms/27.16 ms) were obtained using an inversion recovery sequence (TI=13.14, 29.3, 65.3, 145, 324, 723, 1610, 8000 ms). T2w anatomical imaging scans (TR = 3000 ms, TE_eff_ = 51.26 ms) were obtained with an echo spacing (ESP) of 8.54. ASL and ADC data were acquired using a spin-echo echo-planar imaging (EPI) readout with a field of view (FOV) = 32 x 32 mm², matrix = 64 x 64 mm², thickness = 1 mm, 10 slices. Fast spin-echo multi-slice (FSEMS) scan sequences (TR = 1000 ms, TE_eff_ = 40, ESP = 10 ms) were acquired using a midline and axial orientation with FOV = 50 x 50 mm², matrix = 256 x 256 mm², thickness = 2 mm, single slice. Based on the acquired FSMES-images, the labeling plane for ASL imaging (6.2 mm thickness) was placed in the rat’s neck at a 45° angle to the animal’s rostrocaudal axis. CBF maps were generated by multiphase pseudo-continuous arterial spin labeling (35). ADC maps were generated from diffusion-weighted images acquired in 3 orthogonal directions for b=0/mm² and b=1000 s/mm². DW imaging scans were obtained using a two-dimensional spin-echo echo-planar imaging (SE-EPI) sequence. For contrast-enhanced T1w spin echo, multi-slice images were taken before and immediately after 0.15 ml injection of gadolinium-based contrast agent (GBCA).

#### MRI analysis

Imaging segmentation and image analysis were performed using itk SNAP (Version 3.8.0, 2019) (36). On T2w maps, areas of ischemia were identified as hyperintense regions (37). On T1w maps, the image before Gd injection was subtracted from the image taken after, and areas of BBB breakdown were identified as hyperintense regions (38). Infarct volume was corrected for edema formation using the formula by Kaplan et al. (39): Corrected infarct size = infarct volume x (volume of contralateral hemisphere/volume of the ipsilateral hemisphere). On ADC maps, areas of ischemia were identified as regions of reduced diffusion of at least 23% relative to the same area on the contralateral hemisphere. Similarly, on ASL maps, areas of hypoperfusion were defined as areas with reduced blood flow of at least 57% relative to the same area on the contralateral hemisphere (40). Relative diffusion and perfusion of the pre-defined cortical area were calculated as follows: diffusion/perfusion of the ipsilateral ROI/diffusion/perfusion of the mirrored region on the contralateral hemisphere. In all imaging modalities, ROIs were selected manually by the main investigator blinded to treatment or control (AMS).

### Western blotting

After imaging, rats were killed by intraperitoneal pentobarbital injection (800 mg/kg) (41). Brains were collected and sliced into 2 mm thick coronal sections with an ice-cold stainless-steel matrix (Kent Scientific). Areas of the striatum and cortex of 1 mm^2^ from both hemispheres were taken from one section and snap-frozen on dry ice. Proteins were extracted from the cortical ipsilateral lesion side by cellular lysis using RIPA buffer supplemented with a protease inhibitor cocktail. Protein quantification was performed using the BCA assay (Pierce BCA Protein Assay Kit, 23225, Thermo Fisher Scientific), 50 µg of protein was denatured (95°C for 5 minutes) using lysis and Laemmli sample buffer (161-0737, Bio-Rad, with DTT). Samples were separated on a 10 % gradient gel (Criterion TGX Precast Gel, 5671033, Bio-Rad) using an Electrophoresis Unit (Bio-Rad, Cressier, Switzerland). Proteins were then transferred onto a polyvinylidene difluoride (PVDF) membrane using the electrophoresis unit. Nonspecific binding sites were blocked with PBS containing 5% BSA and 0.1% Tween-20 for 1 hour at room temperature before being incubated in primary antibody (mTOR and phospho-mTOR (Ser 2448), both from Cell Signaling) in 5% PBS-T solution overnight. After 3 x 5-minute-washes in PBS-T, the membrane was incubated in secondary antibody goat anti-rabbit (IgG H&L (HRP), ab6721, Abcam, dilution 1:2000) in 5% BSA PBS-T for 1 hour at RT. After 3 x 5-minute-washes in PBS-T, the membrane was developed using enhanced chemiluminescence (ECL; 12644055, Fisher Scientific) for 5 minutes and immediately imaged after that. Western Blot analysis and quantification were performed using densitometry and corrected for loading using β-tubulin which was used as the housekeeping protein. The membrane was probed for antibodies twice, using the following mild antibody-stripping protocol: The membrane strips were incubated in mild stripping solution (7.5 g glycine, 0.5 g SDS, 5 ml Tween-20, pH to 2.2, and diluted up to 500 ml distilled water) for 2 x 7 minutes, following by 2 x 10-minute-washes in PBS, and 2 x 5-minute washes in PBS-T, before blocking again in 5% BSA and 0.1% Tween-20.

### Statistical analysis

The experimental design and number of animals have been determined and optimized to test the specific hypotheses of the project and follow the standards for scientific reporting and reproducibility (42). Statistical analyses were performed at the end of the experiment, and no further animals were included at that point.

We planned the study with n=9/group, based on the maximum effect size (difference between rapamycin and vehicle) and variability of the change in lesion volume which was defined as the primary outcome (30% with a standard deviation of 20%). We were able to reject the null hypothesis that rapamycin does not change lesion volume with a probability (power) of 0.80. The type I error probability of this null hypothesis (alpha) was 0.05.

Statistical tests were performed using GraphPad Prism 8.42 (La Jolla, USA). The D’Agostino and Pearson normality tests were performed on all data. Appropriate statistical tests were chosen based on the normality of the data. To compare differences between the two treatment groups, an unpaired Student’s t-test was used. The correlation of imaging parameters with functional outcomes was analyzed by simple linear correlation calculation. The correlations were classified as tiny or < 0.05, very small (0.05 < = r < 0.1), small (0.1< = r < 0.2), medium (0.2< = r < 0.3), large (0.3< = r < 0.4), or very large (r > = 0.4) according to Funder and Ozer (43). A P <0.05 is accepted as statistically significant and data are presented as mean ± standard deviation (SD).

## Results

### Excluded animals

A total of 4 rats were excluded from the study, 3 of which had an insufficient LDF drop during filament insertion and 1 due to filament-induced SAH, which occurred before either the treatment or the vehicle was administered.

### Rapamycin improves immediate post-recanalization blood flow

There was no difference in blood flow measured at the time points immediately before re-canalization/treatment administration in control and rapamycin groups, respectively (rapamycin: - 56.58 ± 7.64 % versus vehicle: 66.16 ± 12.35 % pre-MCAo baseline, P=0.1412, Figure 1A). Rapamycin significantly increased the average MCA territory blood for the first 90 minutes post-recanalisation (rapamycin: −9.609 ± 3.374 versus vehicle: −28.32 ± 6.683% of pre-MCAo baseline CBF, P<0.0001, Figure 1B).

**Figure 1.**
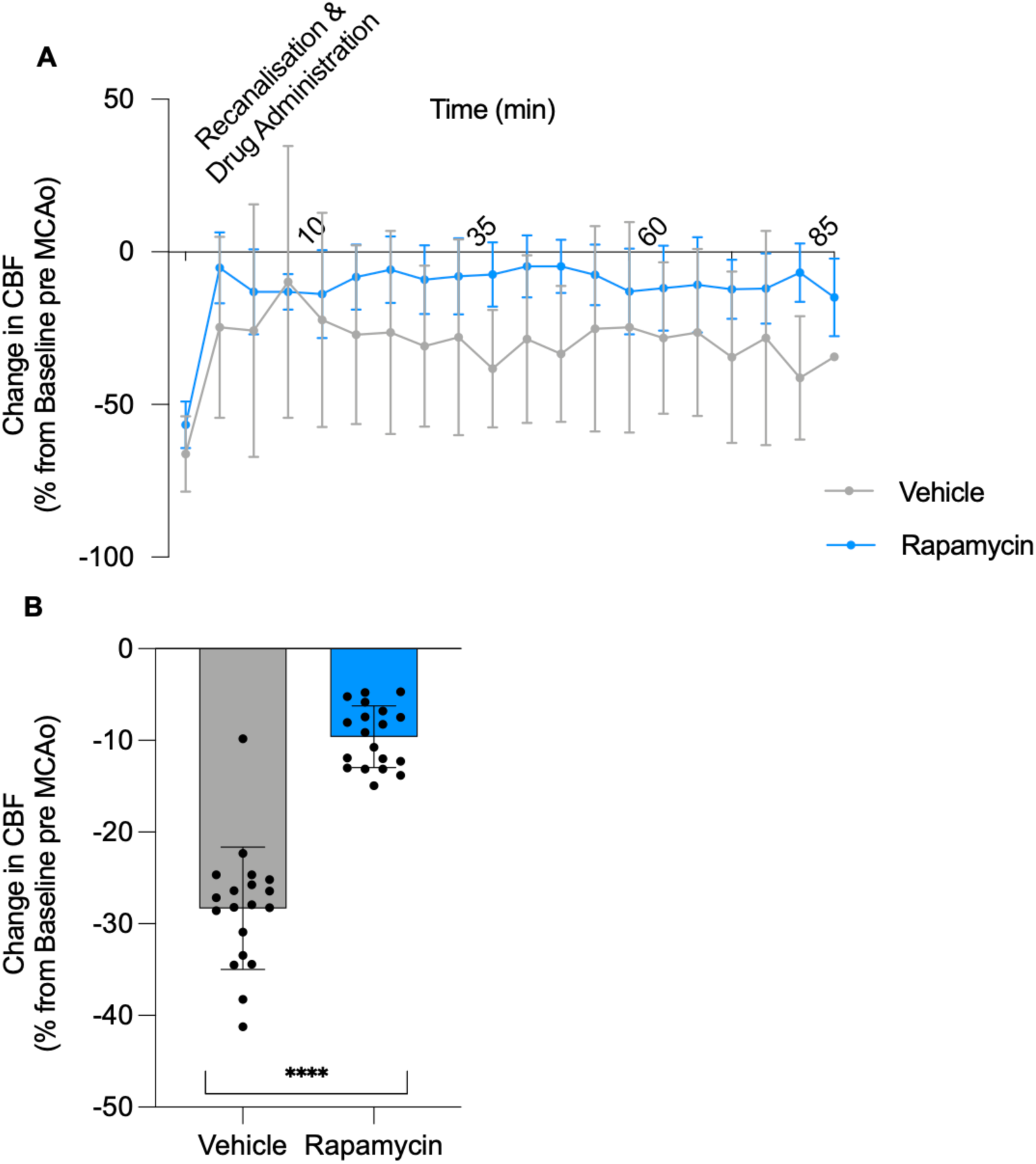
Rapamycin improves post-recanalization blood flow acutely. **(A)** Cerebral blood flow (CBF) of the middle cerebral artery (MCA) territory after recanalization and rapamycin treatment. **(B)** Change in CBF calculated as % change from before middle cerebral artery occlusion (MCAo). Data are mean (SD), ****P<0.0001, compared to vehicle. n=7 for vehicle- and n=9 for rapamycin.

### Rapamycin significantly reduces infarct volume

Rapamycin significantly reduced infarct volume at 3 days post-stroke (rapamycin 44.77 ± 30.93 mm^3^ versus vehicle: 113.3.44 ± 60.19 mm^3^, P=0.0114, Figure 2A). The Waxholm Space Atlas was then overlaid onto the T2w MRI sequences showing that, amongst other areas, the somatosensory cortex was affected by stroke and at least in part salvaged by rapamycin treatment (Figure 2 (B), Supplementary Table 1).

**Figure 2.**
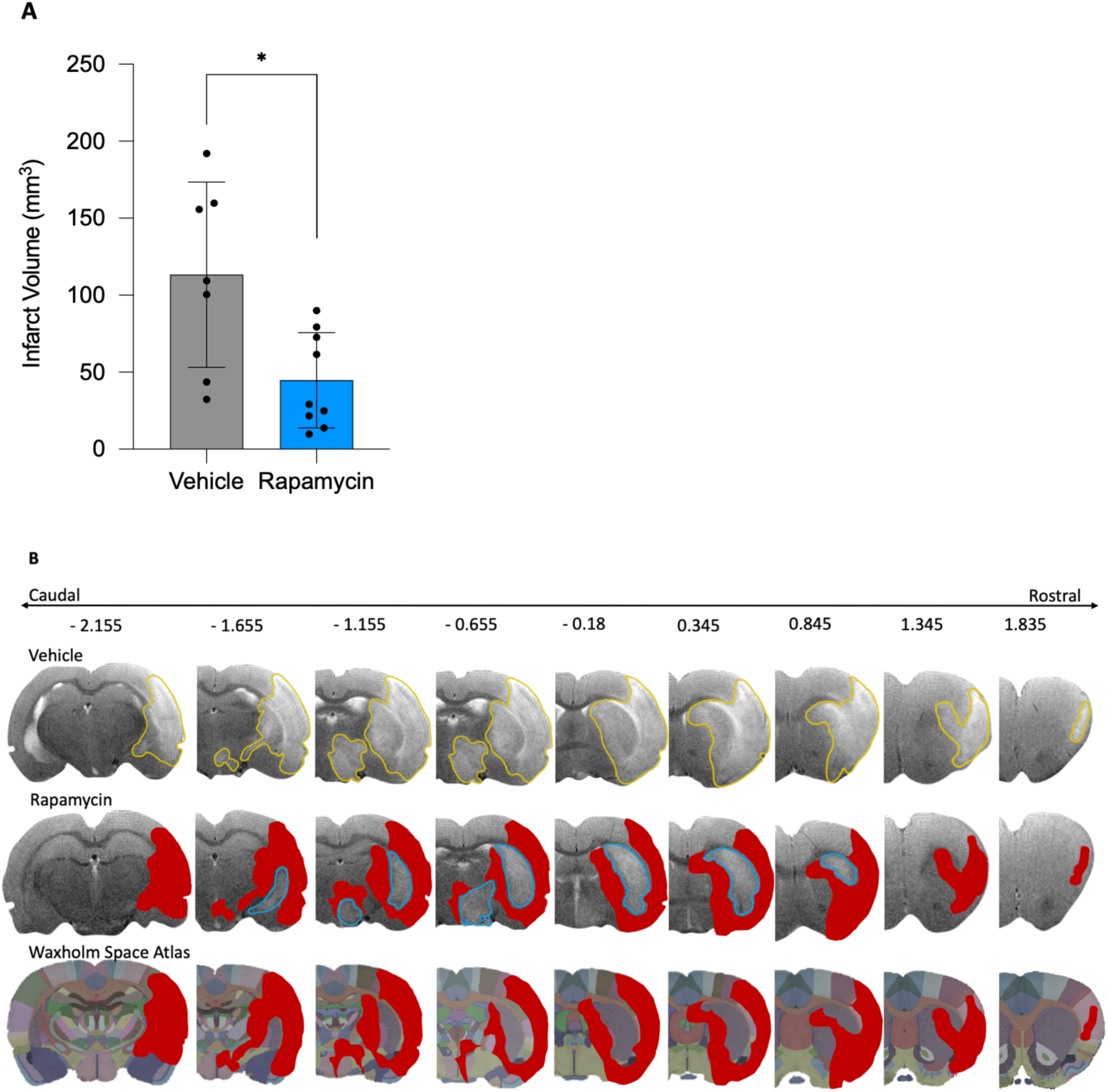
Rapamycin significantly reduces infarct volume. **(A)** Infarct volume assessment with MRI (T2w sequence) 3 days post-stroke, with correction for edema. Data are mean (SD), * P<0.05, compared to vehicle. n=7 for vehicle- and n=9 for rapamycin. **(B)** Coronal view, T2w MRI of representative stroke brains 72 h post tMCAo. Red areas salvaged by rapamycin.

### Rapamycin significantly improves functional outcomes

Rapamycin significantly improved performance on the Garcia test (rapamycin: 12.78 ± 1.039 points versus vehicle: 11.67 ± 0.8660 points, P=0.0295, Figure 3A). Rapamycin did not improve performance on the Bederson test (rapamycin: 1.556 ± 0.7265 points versus vehicle: 2.111 ± 0.6009 points, Figure 3B, P=0.0962). Rapamycin-treated animals required significantly less time to notice the adhesive tape on their forepaw, indicating improved sensory functioning (rapamycin: 45.04 ± 11.91 s versus vehicle: 72.33 ± 12.17 s, P=0.0002, Figure 3 C). Rapamycin-treated animals also required less time to remove the adhesive tape on their forepaw (rapamycin: 49.89 ± 23.09 s versus vehicle: 76.56 ± 12.92 s, P=0.0146, Figure 3D).

**Figure 3.**
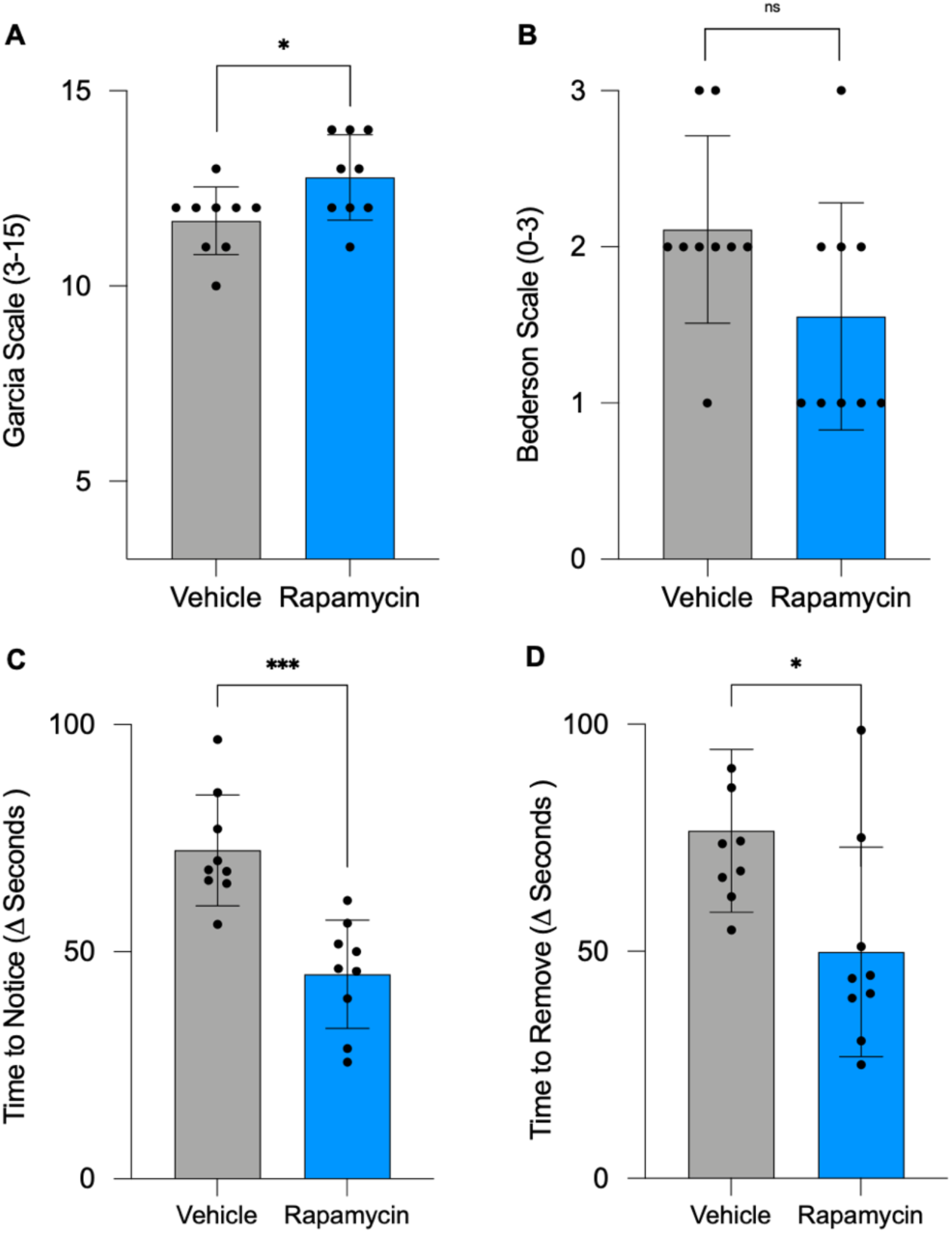
Rapamycin significantly improves functional outcomes. **(A)** Neurological assessment with Garcia scale 72 h post-stroke (3-15 points). **(B)** Neurological assessment with Bederson scale 72 h post-stroke (0-3 points). Data are mean (SD), * P<0.05, when compared to vehicle, n=9 for all groups. **(C)** Rapamycin significantly reduces sensory deficits assessed by the adhesive removal test. Time to notice the left paw (contralateral to stroke side), calculated as the difference between baseline and 72h post-stroke (seconds). **(D)** Rapamycin significantly reduces motor deficits assessed by the adhesive removal test. Time to remove the sticky tape from the left paw (contralateral to stroke side), calculated as the difference between baseline and 72h post-stroke (seconds). Data are mean (SD), *** P<0.001, when compared to vehicle. n=9 for all groups.

There was a tight correlation between infarct volume and neurological test results, with larger lesions leading to worse functional outcomes (Infarct volume vs. Garcia: r=-0.5502, P=0.0272; Figure 4A; Infarct volume vs. Bederson: r=0.6138, P=0.0114, Figure 4B; Infarct volume vs. time to notice of contralateral forepaw in adhesive removal test: r=0.5486, P=0.0278, Figure 4C).

**Figure 4.**
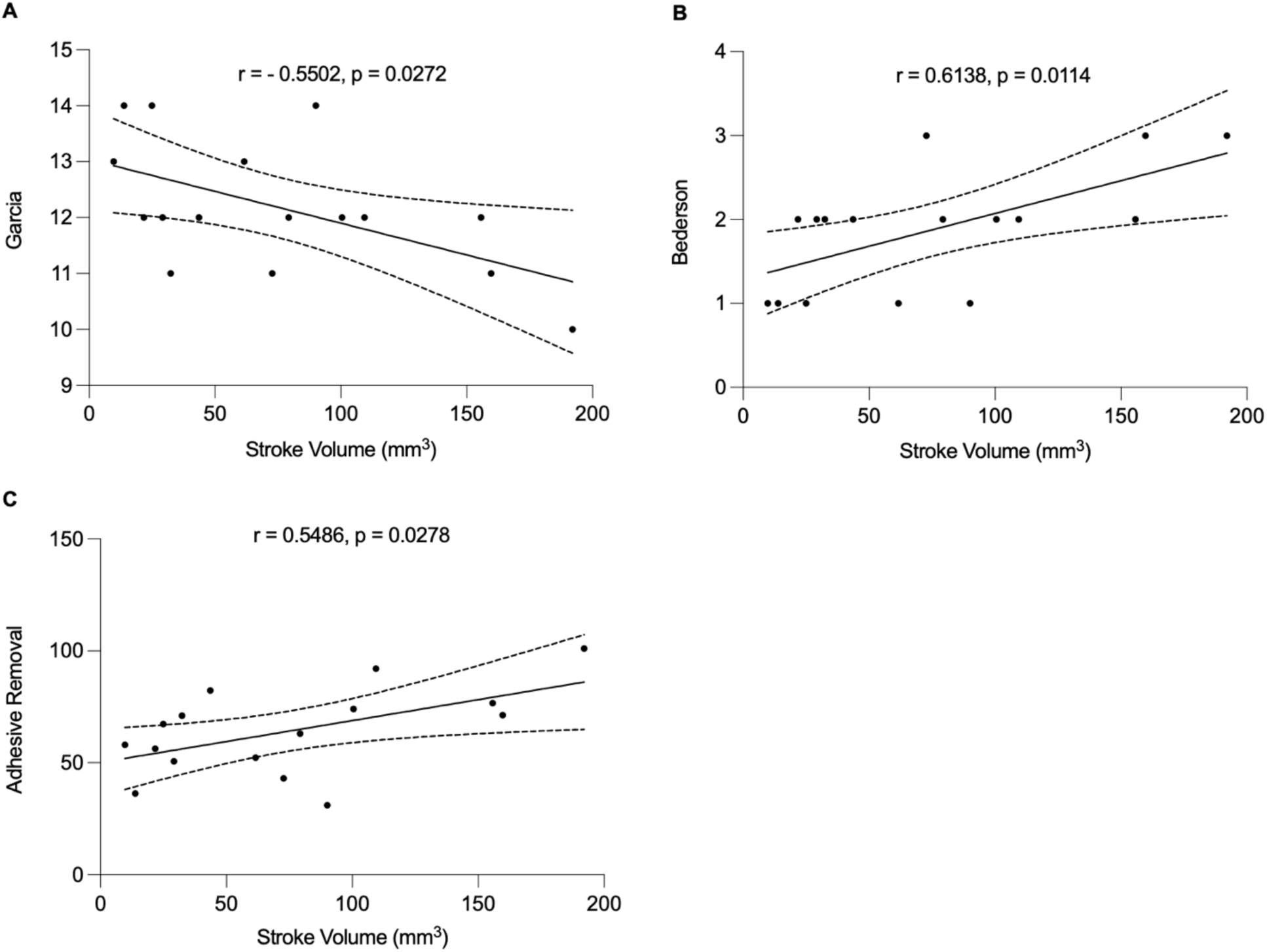
There is a very strong correlation between infarct volume and neurological test results, with larger lesions leading to worse functional outcomes. **(A)** Stroke volume vs. Garcia r=-0.5502, * P<0.05; **(B)** Stroke volume vs. Bederson r=0.6138, * P<0.05; **(C)** Stroke volume vs. time to notice of contralateral forepaw in adhesive removal test r=0.5486, * P<0.05. n=16 for all groups.

### Rapamycin does not alter diffusion or perfusion

After evaluating rapamycin’s effect on blood flow in the hyperacute phase, we next wanted to investigate its effects on spatially matched dynamic changes in the subacute phase of disease progression.

At 3 days, no rat had a large enough diffusion deficit to define the cortical MCA region as ischemic. Rapamycin did not significantly change relative diffusion (rapamycin 1.122 ± 0.04604 versus vehicle 1.109 ± 0.08839, P=0.5765, Figure 5A). Rapamycin did not significantly change absolute diffusion in the ipsilateral ROI rapamycin 0.00081 ± 0.0001 10^4^ mm^2^/sec versus vehicle: 0.00080 ± 0.000 10^4^ mm^2^/sec, P=0.6099, Figure 5B).

**Figure 5.**
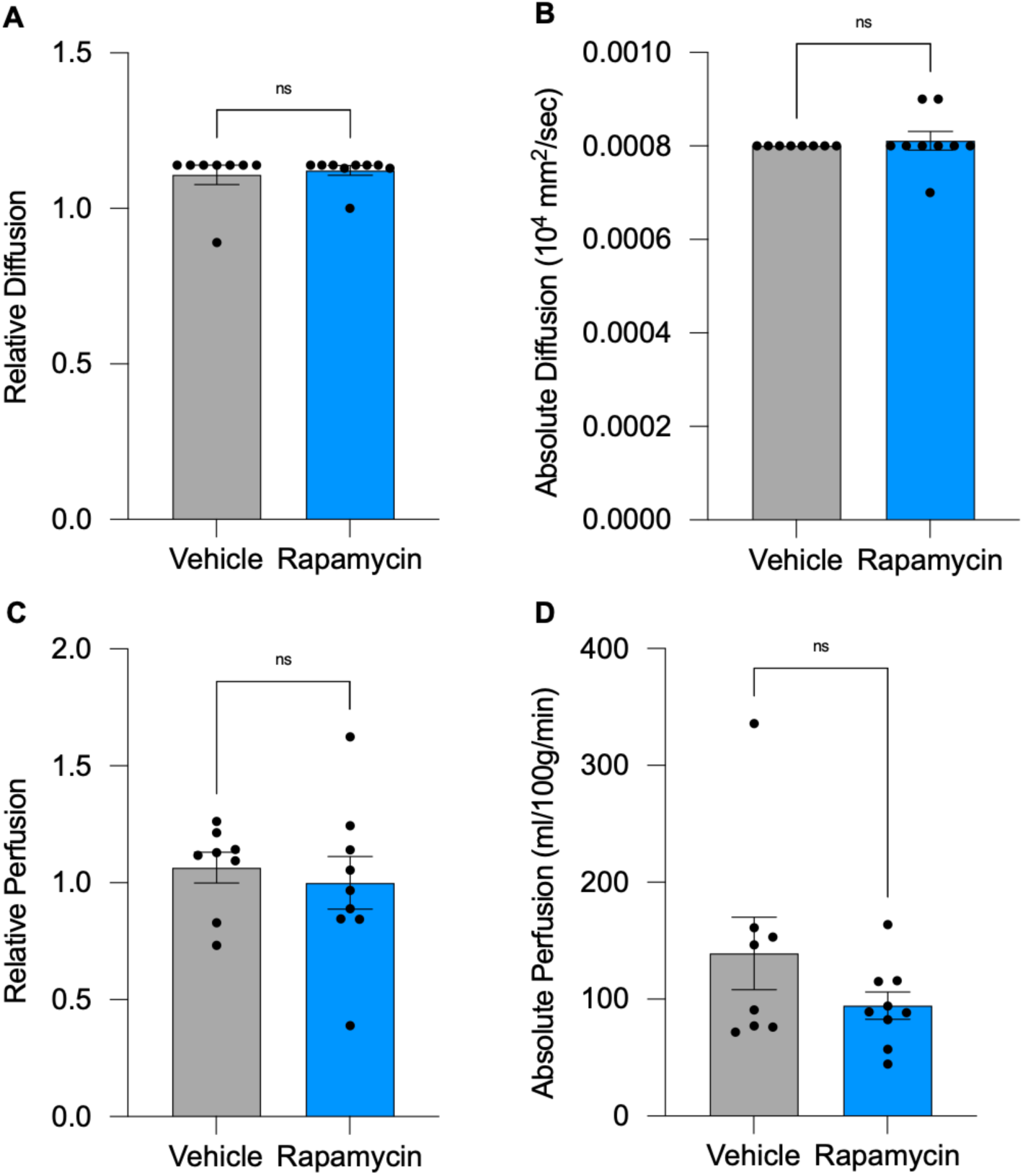
Rapamycin does not alter diffusion or perfusion. **(A)** Ipsilateral diffusion of the stroke area relative to the same area on the contralateral hemisphere was assessed with MRI (ADC sequence) 3 days post-stroke. **(B)** Absolute diffusion values of the ipsilateral hemisphere. **(C)** Ipsilateral perfusion of the stroke area relative to the same area on the contralateral hemisphere assessed with MRI (ASL sequence) 3 days post-stroke **(D)** Absolute perfusion values of the ipsilateral hemisphere. Data are mean (SD), n=9 for rapamycin and n=9 for the vehicle group.

One rat exhibited a stark enough difference with ASL data between the ipsi- and contralateral hemispheres to define a perfusion abnormality. Rapamycin did not significantly change relative perfusion (rapamycin 0.9997 ± 0.3364 versus vehicle: 1.065 ± 0.1856, P=0.6340 Figure 5C). Rapamycin did not significantly change absolute perfusion (rapamycin 94.60 ± 34.6697 ml/100g/min versus vehicle: 139.1 ± 87.92 ml/100g/min, P=0.4807, Figure 5D).

### The effect of rapamycin on edema formation and BBB integrity

Edema formation of the T2w images was assessed using the edema calculation after Kaplan (39): Extent of edema = (volume of the ipsilateral hemisphere – the volume of the contralateral hemisphere)/volume of the contralateral hemisphere. Rapamycin did not significantly change edema volume (rapamycin 9 ± 5.657 mm^3^ versus vehicle: 14.25 ± 7.186 mm^3^, P=0.1129, Figure 6A). Further, T1w imaging with gadolinium injection was performed to assess BBB integrity. 4 additional animals had to be excluded because the amount of Gd reaching the area of interest was insufficient. Rapamycin did not significantly change BBB breakdown (rapamycin 3.270 ± 2.336 mm^3^ versus vehicle: 7.647 ± 4.500 mm^3^, P=0.0606, Figure 6B).

**Figure 6.**
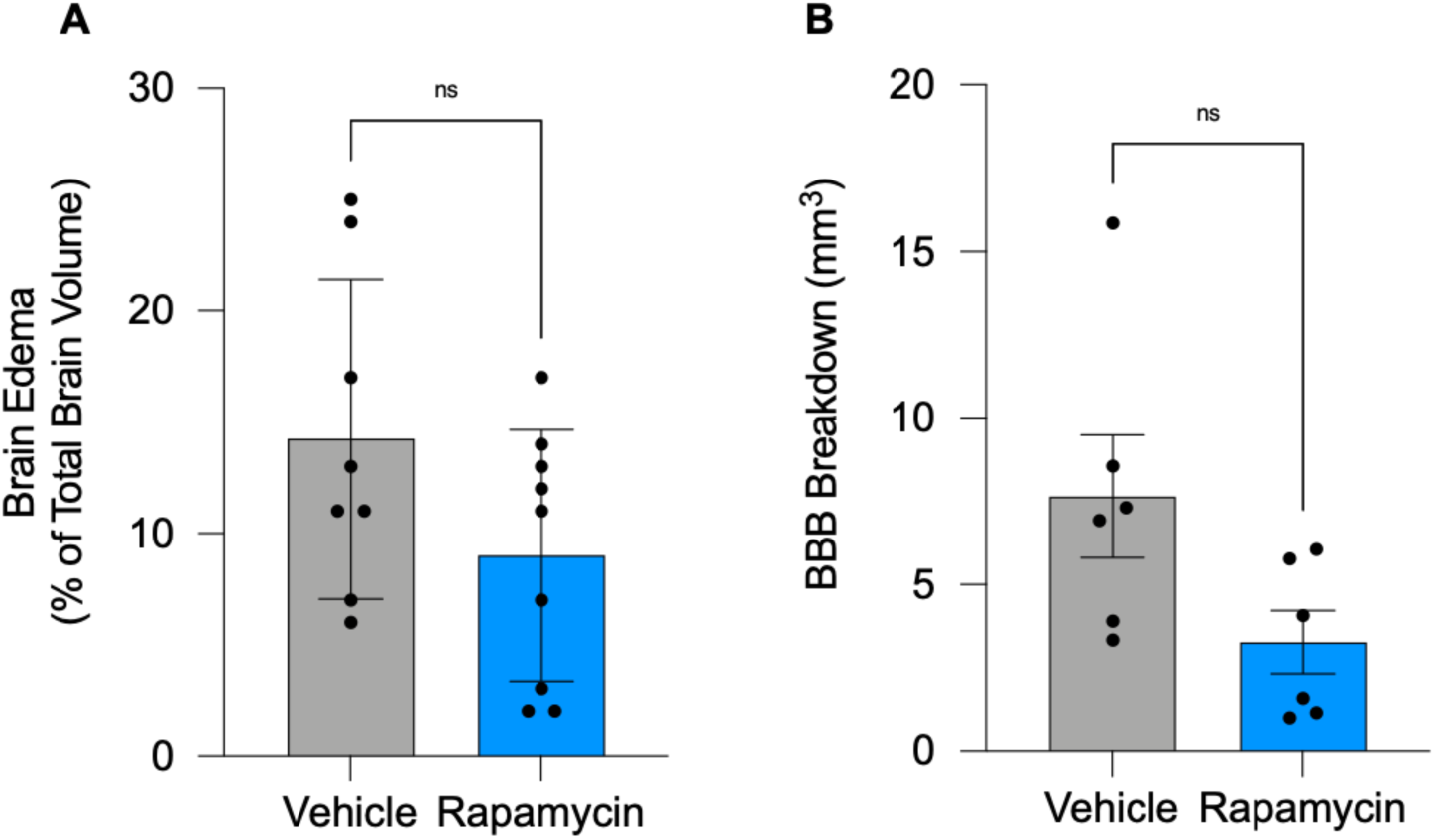
The effect of rapamycin on edema formation and BBB integrity. **(A)** Rapamycin did not reduce edema formation. Brain edema expressed as a percentage of whole-brain volume, assessed with MRI (T2w sequence) 3 days post-stroke. Data are mean (SD), n=8 for control and n=9 for the rapamycin-treated group. **(B)** BBB breakdown assessment with contrast-enhanced T1w spin echo, multi-slice images were taken before and immediately after 0.15 ml injection of gadolinium-based contrast agent (GBCA) and corrected for edema formation. Data are mean (SD), n=6/group in all groups.

### mTOR activity was not reduced at 3 days

For mTOR, the ratio of phosphorylated to total protein can indicate its level of pathway activity, where values over 1 indicate activation and values below 1 suggest an inhibitory effect. The ratio of phosphorylated mTOR (p-mTOR) to total mTOR did not differ significantly in the stroke animals treated with either vehicle or rapamycin (rapamycin: 0.5502 ± 0.3153 versus vehicle: 0.9032 ± 0.4139, P=0.1880, Figure 7).

**Figure 7.**
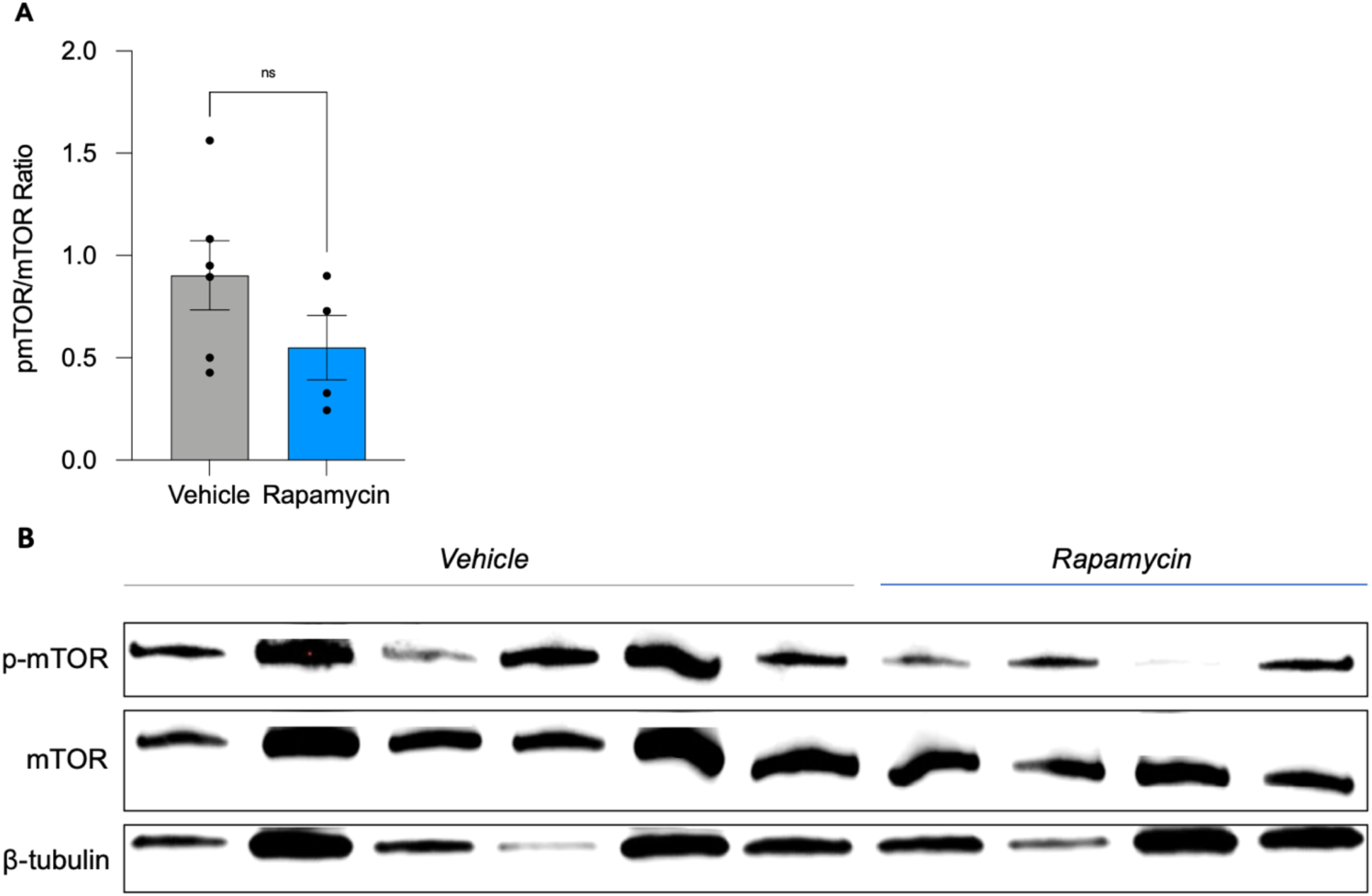
Rapamycin does not inhibit mTOR pathway activity at 3 days. At 3 days, brain tissue from the ipsilateral cortical lesion side from animals treated with rapamycin versus control was used to investigate mTOR activity (measured as pmTOR/mTOR ratio). Western Blot did not show a significant difference in mTOR activity in the lesion area 3 days following the rapamycin treatment. Data are mean (SD), n=4 for rapamycin and n=6 for vehicle.

## Discussion

This study explored the effects of postischemic mTOR inhibition on cerebral blood flow, BBB integrity, and stroke outcome in rats using MRI. The clinically approved mTORC1 inhibitor rapamycin improved reperfusion CBF immediately after recanalization and decreased infarct volume on day 3. Rapamycin also reduced somatosensory deficits in stroke animals and had no effect on BBB, as shown by MRI 3 days following stroke onset.

Rapamycin significantly increased CBF in the MCA territory immediately after recanalizing the occluded artery in rats. This result is in agreement with previous findings from our lab showing increased collateral perfusion and increased post-reperfusion CBF in Wistar and spontaneously hypertensive rats following rapamycin administration at 30 minutes post-stroke (26). Similarly, Wang et al. showed that intraperitoneal rapamycin administration immediately after MCAo in Sprague Dawley rats increased the diameter of deep brain collaterals between the terminal branches of the PCA and MCA (44). Although these findings are encouraging, the timing of administration is of central importance, especially given the potential translational capacity of rapamycin, an already FDA-approved clinically used drug. Therefore, administration of rapamycin after recanalization, as in this study, is of greater clinical significance compared to studies in which the drug was administered before or at stroke induction (9, 13, 14, 18, 19). The mechanism by which rapamycin exerts its beneficial vasodilatory effects is not fully elucidated, but studies by our group and others suggest an endothelial nitric oxide synthase (eNOS) dependent mechanism as seen in mouse models and isolated collateral vessels, a RhoA-dependent pathway, or direct reduction in calcium sensitivity in smooth muscle cells (26, 45–47). Better understanding the pathway by which rapamycin exerts its beneficial effects, which cell types are primarily targeted, and which dose exerts optimal effects are essential questions for future studies.

Rapamycin significantly reduced lesion volume at 3 days when compared to the vehicle-treated group. Our group’s previous meta-analysis on rapamycin treatment in rodent models of stroke included a global estimate of 30 comparisons and suggested that rapamycin significantly reduced infarct volume by 22% (2). The current study exceeded the expected number with a mean reduction of 39%. MRI imaging didn’t show a difference in diffusion or perfusion when rapamycin-treated animals. Further, no diffusion-perfusion mismatch (DPM) was observed in either the treatment or control group, suggesting that there was no “salvageable” brain tissue at this time point, which might also explain why there was no effect in the treatment group. Literature suggests that DPM volume gradually decreases over time, with a stark decrease to a fourth of the initial penumbral region from 45 to 210 minutes after permanent MCA occlusion (27, 48). There have been far fewer studies on the evolution of DPM after recanalization in temporary MCAo. We found one study by Meng et al. that found the DPM was greatest within the first 2 hours following reperfusion (49). Therefore, a two-hour window post reperfusion is thought to have the most salvageable tissue and therefore be the optimal time for the administration of adjunct cytoprotective therapies. It is likely that by administering rapamycin at the time of recanalisation we were most likely to have an effect as there is still brain left to save. This is evidenced by a reduced infarct volume at 3 days.

BBB integrity was not improved in rapamycin-treated animals versus control. Subtle impairment of BBB function has been reported as early as 30 minutes to 6 hours after ischemia-reperfusion (I/R) in a mouse model of t-MCAo, resulting in extravasation of small macromolecules (≤3 kDa) from blood to brain parenchyma (50–52). After more than 3 hours, larger macromolecules (≥ 40kDa) leak through the barrier, contributing to high BBB leakage during this phase. Tight junction maintain their integrity during the hyperacute phase following the insult but lose their integrity at around 48 hours post-I/R (51, 53, 54) due to damage caused by cytokine-activated matrix-metalloproteinases MMP-3, MMP-9, and cyclooxygenase-2. The loss of tight-junction integrity during the acute phase of BBB breakdown is thought to be the leading cause of the development of vasogenic edema (50, 55). Previous studies have reported that rapamycin treatment significantly reduced BBB breakdown after stroke (8, 9, 13, 14, 19, 56). A potential explanation for the lack of benefit of rapamycin treatment is that assessment of BBB breakdown in our study was a secondary analysis and it is possible that we were underpowered to detect an effect. Alternatively, the lack of effect may have been due to the concentration of rapamycin dropping at 3 days (as evidenced by our Western blotting data), the timepoint when BBB breakdown is known to peak following ischaemic stroke. Our findings of a lack of effect of rapamycin on the BBB is also consistent with our T2W brain edema analysis, which also showed that rapamycin did not significantly reduce edema volume at 3 days.

Rapamycin significantly improved somatosensory function after stroke. Studying behavior is very important for evaluating the effectiveness of neuroprotective therapy to understand its impact on functional recovery. Three days is an optimal time point to assess potential long-lasting functional deficits and improvements with treatment, as it is not masked by acute deficits and weaknesses due to the surgery. The testing battery chosen in this study aimed to include both functional and neurological deficit scores that test broad motor and somatosensory control and overall functional outcome. We found that rapamycin significantly improved the time to notice and time to remove the adhesive in the adhesive removal test. Interestingly, time to notice (an assessment of somatosensory function) showed greater improvement in rapamycin treatment compared to the time to remove (encompassing both somatosensory and motor components) (57). Furthermore, time to notice showed a stronger correlation with final infarct volume than time to remove. These differences may be due to the increased sensitivity of time to notice to detect sensory disturbances owing to the somatosensory cortex being affected by MCAo in the rat and potentially subsequently salvaged by rapamycin treatment. This was depicted in Figure 2B and Supplementary Table 1, where the Waxholm Space Atlas was overlaid onto the T2w MRI sequences showing that, amongst other areas, the somatosensory cortex was affected by stroke and at least in part salvaged by rapamycin treatment.

Alternatively, these findings could simply be due to the stroke surgery itself requiring the ligation of the external carotid artery thus inducing ischaemia of muscle of mastication making it more difficult for the animal to remove the adhesive with its mouth (58). Rapamycin treatment did not significantly improve the outcome of the Bederson or Garcia tests, which assess sensorimotor deficits, independent of the use of the mouth (59). Therefore our results suggest that rapamycin was better at rescuing sensory deficits compared to motor deficits in our model. Our findings are in agreement with previous studies that showed that rapamycin treatment improved neurological outcomes (5, 6, 10, 11, 13, 14, 19, 60). Our previous study reported that rapamycin administered at the time of stroke significantly improved the time to notice on the adhesive removal test (26). Furthermore, our recent systematic review and meta-analysis of rapamycin in stroke also found that rapamycin improved neurological scores by 30% and there was a significant correlation between infarct volume and neurobehavioural scores (2).

Western blot of tissue from the ipsilateral cortical lesion showed that mTOR activity (as measured by pmTOR:mTOR ratio) at three days was not downregulated at 3 days post-stroke. suggesting that rapamycin had already exerted the entirety of its effects. To the best of our knowledge, there is no information on mTOR activity after rapamycin treatment in rats after 3 days. However, a study in Sprague Dawley rats showed that 72 hours following a 1 mg/kg dose of rapamycin, the blood concentration of rapamycin dropped to ∼1 nM (estimated half-life of 25 hours), which is below the effective concentration needed for mTOR inhibition in neuronal cell culture (61, 62). Another study by Nalbandian et al reports that rapamycin’s half-life is 58 hours in mice (63). Conversely, Böttiger and colleagues reported rapamycin’s half-life to be 79 hours in healthy males, suggesting a species-dependent difference in the biochemical effect and metabolism of the drug (64). To test this, future studies should repeat the experiments but choose earlier study endpoints and compare the effects of rapamycin treatment on mTOR activity, the microvasculature, and infarct volume with MRI, histology, and functional outcome. Alternatively, to test whether the positive effects of rapamycin treatment are due to its mediating effects on the blood flow in the hyperacute phase (i.e. within the 2-hour window of salvageable tissue post-recanalisation), treatment could be started after 2 hours post-recanalisation to test whether the beneficial effect of rapamycin is maintained, diminished or lost. A better understanding of the temporal pattern under which post-ischemic mTOR inhibition exerts its beneficial effects on stroke outcomes might increase our knowledge of the underlying physiological properties of rapamycin in the context of stroke and eventually optimize the treatment approach for a clinical setting.

In conclusion, this is the first high-quality preclinical trial of rapamycin taking into account key translational considerations. We found that inhibiting mTOR activity upon reperfusion with the FDA-approved drug rapamycin in an ischemic rat model immediately increases CBF. At 3 days, lesion volume was significantly smaller, and functional outcome improved. Thus our study confirms previous findings of rapamycin’s positive impact on stroke outcomes, reinforcing its potential as a cerebroprotective drug for ischemic stroke patients. Confirmation of our positive results in rodents using larger animal models is an important next step for establishing the translatability of our research findings.

## Acknolwedgements

AMB is currently funded by a Trans-Atlantic network grant from the Leducq Foundation and by an Einstein Visiting Fellowship to the Charite in Berlin, from the Einstein Foundation, Berlin. YC is funded by Alzheimer’s Research UK. DJB was funded by the Medical Research Council UK (MR/M022757/1) and the Australian National Health and Medical Research Council (APP1182153).

## Author Contributions

AMS, YC, JL, AMB, and DJB contributed to the study conception and design. Material preparation, data collection, and analysis were performed by AMS, YC, JL, and DJB. The first draft of the manuscript was written by AMS, and all authors commented on previous versions of the manuscript. All authors read and approved the final manuscript.

## Disclosures/conflicts of interest

AMB is senior medical science advisor and co-founder of Brainomix, a company that develops electronic ASPECTS (e-ASPECTS). The other authors declare no competing conflict of interest.

## Supplementary information available on JCBFM Website

