## Supplemental table 1 for "Post-stroke rapamycin treatment improves post-recanalization cerebral blood flow and outcome in rats"

**Supplementary data**

**Table 1**. Area affected by MCA occlusion and salvaged by rapamycin. Based on visual representation overlay of T2w MRI images onto Waxholm Space Atlas of Sprague Dawley Rat Brains (65).

| **AP (mm)** | Area affected by MCA occlusion (bold areas salvaged by rapamycin) |
| --- | --- |
| 1.835 | - **Primary somatosensory area, face representation** - **Granular insular cortex** |
| 1.345 | - **Primary somatosensory area, face representation** - **Secondary somatosensory area** - **Insular cortex** - **Piriform cortex** - **Claustrum** - **Caudate putamen** |
| 0.845 | - **Primary somatosensory area (forelimb representation, dysgranular zone, face representation)** - **Secondary somatosensory area** - **Insular cortex** - **Piriform cortex** - **Claustrum** - **Caudate putamen** |
| 0.345 | - **Primary somatosensory area (forelimb representation, dysgranular zone, face representation)** - **Secondary somatosensory area** - **Insular cortex** - **Piriform cortex** - **Claustrum** - Caudate putamen |
| - 0.18 | - **Primary somatosensory area (forelimb representation, dysgranular zone, barrel field, face representation)** - **Secondary somatosensory area** - **Insular cortex** - **Piriform cortex** - **Claustrum** - Caudate putamen |
| - 0.655 | - **Primary somatosensory area (forelimb representation, dysgranular zone, barrel field)** - **Secondary somatosensory area** - **Insular cortex** - **Piriform cortex** - Caudate putamen - Basal forelimb region, unspecified - **Hypothalamic region, unspecified** |
| - 1.155 | - **Primary somatosensory area (forelimb representation, dysgranular zone, barrel field)** - **Secondary somatosensory area** - **Insular cortex** - **Piriform cortex** - Caudate putamen - **Basal forelimb region, unspecified** - **Hypothalamic region, unspecified** |
| - 1.655 | - **Primary somatosensory area (dysgranular zone, barrel field)** - **Secondary somatosensory area** - **Insular cortex** - **Piriform cortex** - Caudate putamen - **Globus pallidus external (lateral part)** - **Corticofugal tract and corona radiata** - **Basal forelimb region, unspecified** - **Hypothalamic region, unspecified** |
| - 2.155 | - **Primary somatosensory area (dysgranular zone, barrel field)** - **Secondary somatosensory area** - **Insular cortex** - **Piriform cortex** - **Caudate putamen** - **Globus pallidus external (lateral part)** - **Corticofugal tract and corona radiata** |
